# Signaling activated by nucleolar localised Notch4 Intracellular Domain underlies protection from genomic damage

**DOI:** 10.1101/670588

**Authors:** Neetu Saini, Apurva Sarin

**Affiliations:** Institute for Stem Cell Science & Regenerative Medicine (inStem), Bellary Road, Bengaluru, Karnataka, India; Department of Biology, Manipal Academy of Higher Education, Manipal, India

**Keywords:** Notch4, Nucleolus, Apoptosis, Genomic damage, Ribosomal biogenesis

## Abstract

The assembly of signaling hierarchies and their spatiotemporal organization together, contribute to diverse signaling outcomes. This is evident in the Notch pathway, which regulates an array of cellular processes, despite a small number of core components. Here, we describe a Notch4 activated signaling cascade, dependent on the nucleolar localization of the Notch4 Intracellular Domain (NIC4), that protects cells from genotoxic damage. Localization was assessed by immune-staining for endogenous Notch4 and visualization by confocal microscopy, in breast cancer cell lines. Live-cell, imaging-based, biophysical analysis of NIC4-GFP expressing cells, indicated unhindered mobility between the nucleolus and nucleoplasm and a stable nucleolar pool of NIC4-GFP. RNAi-mediated ablations, coupled with analysis of recombinant forms of NIC4 with modifications of its nucleolar localization sequence, confirmed nucleolar localization and identified the nucleolar proteins, Nucleolin and Fibrillarin, as key intermediates in the NIC4-activated signaling cascade. The transcriptional control of ribosome biogenesis (47s and 45s pre-rRNA transcription), emerged as another unexpected consequence of the subcellular distribution of NIC4. Taken together, this study describes intrinsic features of NIC4 that confer spatial flexibility and expand the repertoire of Notch4 signaling.

## Introduction

Signaling through the Notch receptors protects from diverse apoptotic stimuli [1,2,3,4]. In mammals, four Notch receptors Notch1-4 are known, which are activated by binding to one of five ligands, Delta-like 1/3/4 or Jagged 1/2 [5,6]. Notch receptors typically contain an extracellular domain consisting of multiple EGF-like repeats, a trans-membrane domain and an intracellular domain [7]. Upon ligand binding, the Notch receptor undergoes a series of proteolytic cleavages, releasing the intracellular domain (NIC) into the cytoplasm [8,9,10]. In the nucleus, NIC complexes with cofactors RBPj-κ and Mastermind like (MAML) to form an activation complex, which induces transcription of various key genes [11,12].

Apart from this core canonical pathway, there are several reports of atypical ligand-independent and non-nuclear Notch signaling in diverse systems [13,14,15,16,17]. While Notch1 signaling has been explored in multiple development and disease contexts; studies investigating Notch4 have mainly focused on its role in the endothelial system and resistance to treatment in cancers [3,18,19,20]. Hence, the molecular regulation of Notch4 signaling is not completely elucidated. Here, the intra-cellular response to agents that trigger genomic damage is used to characterize the mechanism underlying one aspect of NIC4-mediated signaling in mammalian cells. Building upon its observed distribution in the nucleolus (and nucleoplasm), NIC4-mediated anti-apoptotic activity is assessed for dependence on nucleolar proteins, canonical regulators of NIC transcription, as well as molecules implicated in sensing and repair of genomic damage. The analysis of molecular dynamics and spatially restricted recombinant forms of NIC4, revealing specific functions of nucleolar pools of NIC4 are described. Apart from demonstrating a requirement for nucleolar proteins in protection from genomic damage, these experiments uncover an unexpected role for NIC4 in ribosome biogenesis. Taken together, the data show that spatial regulation i.e. its nucleolar localization, expands the repertoire of signaling cascades activated by NIC4.

## Materials and Methods

### Cells

HEK 293T (HEK), MDA-MB-231 cell lines were obtained from ATCC (Manassas, VA, USA), Hs578T, BT-459 and SUM149 were obtained from TR Santhosh Kumar (Rajiv Gandhi Centre for Biotechnology (RGCB), Thiruvanthapuram, India); the HCC1806 cell line was obtained from A. Rangarajan (Indian Institute of Science, Bengaluru, India) and MCF-7 cells were from D. Notani (National Centre for Biological Sciences, Bengaluru, India). HEK293T and MDA-MB-231 cells were maintained in DMEM (GIBCO, Life Technologies USA) supplemented with 0.1% penicillin/streptomycin and 10% FBS (Scientific Hyclone TM, Waltham, MA, USA) at 37°C with 5% CO_2_. HCC1806, BT-549, Hs578T and SUM149 cells were maintained in RPMI-1640 supplemented as above.

### Reagents

5-Fluorouracil (F6627), 4-Nitroquinoline *N*-oxide (N8141) and Thapsigargin (T9033) were from Sigma-Aldrich (St. Louis, MO, USA). Etoposide (341205) was from Calbiochem-Merck Millipore (Darmstadt, Germany). Trizol and Superscript First Strand Synthesis System were from Invitrogen. SYBR™ Green Master Mix was from Thermo Scientific (CA, USA). Dharmafect-1 and siRNA to the scrambled control (D-0018010-10), Notch4 (L-011883-00), Notch1 (L-007771-00), RBPj-κ (L-007772), Fibrillarin (L-011269), Nucleolin (L-003854), Rad50 (L-005232) and Nbs1 (L-009641) were from Dharmacon (Lafayette, CO, USA). Antibodies to Notch4 (2423) and Nucleolin (14574) and anti-rabbit Alexa 543 were from Cell Signaling Technology (MA, USA). All other reagents were purchased from Invitrogen (CA, USA).

### Plasmids

Human NIC4 was sub-cloned into pEGFP-N3 (BD Clontech, Mountain View, CA) between EcoRI and BamHI restriction sites to obtain NIC4-GFP using the following primers:

NIC4- EcoRI Forward: 5’-ATAGAATTCAATGCGGCGTCGAC-3’

NIC4-BamHI Reverse:5’-TTAGGATCCTTTTTTACCCTCTC-3’

NoLS_NIC 4 and NIC43RA mutants were prepared using PCR mediated mutations and addition of NIK (RKKRKKK) NoLS signal sequence to the former using the following primers :

NoLS_NIC4 Forward: 5’-TAGAATTCATGCGGAAGAAACGGAAGAAGAAGCGGCGTCGACGCCGAG-3’

NoLS NIC 4 Reverse: 5’-AATGGATCCTTTTTTACCCTCTCCTCCTTG-3’

The following primers were used for the generation of the NIC43RA-GFP construct using PCR based site directed mutagenesis:

NIC43RAForward: 5’-GCGCCTGCGACTCAGTCAGCTCCCCACCGACGCGC GCCCCCACTAGGCGAGGACAGC-3’

NIC43RAReverse: 5’-CGCGCGTCGGTGGGGAGCTGACTGAGTCGCAGGCG CTCGAGTGAAACCAGGGGGCAGC-3’

mTagRFP-T-Fibrillarin-7 was a gift from Michael Davidson (Addgene plasmid # 58016); GFP-Nucleolin from Michael Kastan (Addgene plasmid # 28176) and Human Bcl-xL GFP plasmid from Richard J. Youle (National Institutes of Health, Bethesda, MD). Construct sequences were verified by automated Sanger sequencing conducted in-house.

### Transfections

HEK cells grown in flasks were trypsinized and seeded at a density of 0.25 × 10^6^ cells in (tissue culture grade) 35mm culture dishes (Greiner Bio-one, Kremsmünster, Austria). 100nM siRNA or plasmids at indicated concentrations were transfected with Dharmafect or Liofectamine 2000 as per the manufacturer’s instructions when cultures were 50-60% confluent (by 24 h post-plating). Cells transfected with siRNA were incubated for 24-26 h before being harvested by trypsinization and re-plated before plasmid transfections (with Lipofectamine 2000), 24 h after the last re-plating. Twenty-four hours after transfection, cells were treated with chemicals as described below. MDA-MB-231 and Hs578T cells were plated at 0.05 - 0.06 ×10^6^ cells per well in wells of a 24-well plate for transfections the next day at 60-70% confluency. siRNA was transfected using RNAi MAX (Invitrogen, USA), following the manufacturer’s instructions. Plasmids were transfected using Lipofectamine 2000 or Lipofectamine LTX at the following concentrations: NIC4-GFP (2µg), pEGFP-N3 (1µg), Bcl-xL GFP (2µg); in MDA-MB-231 cells, NIC4-GFP (1.5µg), pEGFP-N3 (0.5µg). Total DNA transfected in different transfection groups were equalized with pcDNA3. Silencing was estimated by analyzing transcript levels of the genes in cells transfected with control or gene-specific siRNA. 48 h post siRNA transfection, 0.5×10^6^ cells were lysed using TRIzol (Invitrogen) followed by RNA isolation as per manufacturer’s instructions. 2μg of RNA was used for cDNA synthesis using Superscript First Strand cDNA synthesis kit and real time PCR was setup using gene specific primers.

### Induction of apoptosis and assays for cellular damage

To assess apoptotic damage, 24 h post-transfection cells were treated with Etoposide (10µM) or 5-Fluorouracil (10µM) or 4-Nitroquinoline *N*-oxide (5µM), for 48 h in serum free DMEM (HEK293T) or 2.5% serum containing DMEM (MDA-MB-231). Cells were treated with Thapsigargin (10µM) for 20 h in serum-free DMEM. Cells were harvested and stained with Hoechst 33342 (1μg/ml) and cells scored for nuclear damage in GFP positive cells using a fluorescent microscope (Olympus BX-60). Approximately 200 cells in five random fields were scored for apoptotic damage.

### Immunostaining

For immunostaining Notch4, Notch1 and Nucleolin, the following protocol was used. Cells were plated at 0.3 ×10^6^ cells in cut confocal dishes and cultured for 48 h to adhere and increase in number. The monolayer was fixed with 2% PFA (freshly reconstituted) and incubated in the dark for 20 min at ambient temperature. Dishes were permeabilized using 0.2% Triton-X 100 for 10 min at ambient temperature and blocked in freshly made buffer (5% Normal goat serum, 0.3% Triton) for one hour at ambient temperature. Samples were treated with primary antibody, added at the indicated dilutions in 5% BSA-PBS, Nucleolin (1:100), N4 (1:100), N1 (1:100), for 2.5 h at ambient temperature. Samples were washed 2x in PBS and secondary fluorescence-conjugated antibody was added and samples incubated for 1 h, protected from light at ambient temperature. Samples were washed 2x with PBS, counterstained with Hoechst 33342 and imaged.

### Real Time-PCR

Total RNA was isolated using TRIzol reagent according to manufacturer’s instructions and concentration was determined using the Nanodrop 2000 (Thermo Scientific). 2µg of total RNA was used for cDNA synthesis using SuperScript First-Strand Synthesis System. Real-time PCR was performed using Maxima™ SYBR Green qPCR Master Mix and Bio-Rad CFX96 Touch™ Real-Time PCR Detection System. Relative change in gene expression was calculated using 2–ΔΔCt method using as GAPDH as reference gene [21]. The primers used for RT PCR are as follows:

Gene Forward (5’-3’) Reverse (5’-3’)

GAPDH: TGCACCACCAACTGCTTAGC; GGCATGGACTGTGGTCATGAG

FBL: TGGACCAGATCCACATCAAA; GACTAGACCATCCGGACCAA

NCL: CCAGCCATCCAAAACTCTGT; TAACTATCCTTGCCCGAACG

RAD50: GGGTTTCCAAGGCTGTGCTA; TCTGACGTACCTGCCGAAGT

NBN: CACTCACCTTGTCATGGTATCAG; CTGCTTCTTGGACTCAACTGC

HES5: CCGGTGGTGGAGAAGATGCG; GCGACGAAGGCTTTGCTGTG

HES1: AGGCTGGAGAGGCGGCTAAG; TGGAAGGTGACACTGCGTTGG

RBPJK: AACAAATGGAACGCGATGGTT; GGCTGTGCAATAGTTCTTTCCTT

Notch4: GCGGAGGCAGGGTCTCAACGGATG; AGGAGGCGGGATCGGAATGT

Notch1: TCCACCAGTTTGAATGGTCA; AGCTCATCATCTGGGACAGG

47SrRNA: TGTCAGGCGTTCTCGTCTC; AGCACGACGTCACCACATC

45SrRNA: GCCTTCTCTAGCGATCTGAGAG; CCATAACGGAGGCAGAGACA

### Co-localization analysis

HEK cells transfected with NIC4-GFP and Fibrillarin-RFP (1µg), were cultured for 24 h post-transfection and fixed with 2% PFA. Images were acquired using Olympus FV3000 confocal microscope (63 X NA 1.35 oil-immersion objectives). Co-localization of NIC1-GFP or NIC4-GFP with Fibrillarin-RFP was quantitated using co-localization threshold plugin after removing background in Fiji Image J software. Manders correlations coefficient is proportional to the fraction of fluorescence intensity of one channel that co-localizes with the other channel and ranges from 0 (no co-localization) to 1 (maximum co-localization).

### FRAP and FLIP analysis

HEK cells were co-transfected with NIC4-GFP (0.5µg) and Fibrillarin-RFP (0.5µg), 18-24 h prior to analysis. For these experiments, cells were plated and transfected on sterile cover-slips fixed in Petri-dishes to allow for confocal imaging and analysis of cells without trypsinization. Cells were imaged using Olympus FV3000 (oil immersion objective, 63X. 1.35 NA). The FRAP module was used to photo-bleach and acquire images every 2 sec immediately after photo bleaching the region of interest (ROI, white arrowhead in images). Fluorescence recovery was quantified after correcting for photo-bleaching using cellSens software.

For the Fluorescence Loss In Photobleaching (FLIP) analysis, HEK cells were transfected with NIC4-GFP (0.5µg) and Fibrillarin-RFP (0.5µg) using Fugene HD and cultured for 24 h. FLIP analysis was performed on Olympus FV3000 at 60x oil objective with 37 °C stage incubator. ROI was drawn around nucleus excluding nucleolus and photo-bleaching (of ROI) was done for 700 milliseconds with 60% of laser (488 or 561). Time lapse images were acquired every 2 sec before and after bleaching. Fluorescence intensities of NIC4-GFP and Fibrillarin-RFP in the ROI restricted to the nucleolus were quantified using Fiji Image J software.

### Statistical Analysis

Data are represented as mean ± standard deviation (Mean ± SD) for three independent experiments. Statistical significance was measured using two-tailed Student’s t-test and p values < 0.05 were considered to be statistically significant. Data plotted for FRAP and FLIP analysis are mean ± SD from two experiments

## Results

### Notch4 regulates susceptibility to genomic damage

Chemicals such as etoposide and 5-fluorouracil (5-FU) target the genome triggering damage to DNA and apoptosis in mammalian cells [22,23]. Implicating Notch signaling in protection from genomic damage, RNAi-mediated ablation of Notch4 increased the sensitivity of the breast cancer cell line MDA-MB-231 to apoptosis triggered by etoposide or 5-FU (Fig. 1A). Conversely, expressing the processed intracellular domain of Notch4 (NIC4), protects cells from apoptotic damage (Fig. 1B). Immuno-staining and visualization by confocal microscopy of its subcellular distribution, revealed discrete intense areas and a more generalized or diffuse distribution in the nucleus (green, Fig. 1C), as well as staining at the cell membrane (Fig. 1C and Supplementary Figure 1A). Intriguingly, the brighter intense spots overlapped with staining for Nucleolin (red, Fig. 1C), indicating co-localization at the nucleolus [24]. Notch4 staining was not detected in the nucleus in cells treated with a gamma-secretase inhibitor (GSI)-X, which block Notch processing at the membrane and release of the intracellular domain (Fig. 1D and Supplementary Figure 1B), indicating that the antibody detects processed Notch4 (NIC4). The non-nuclear staining seen in GSI-X treated groups, is most likely reporting full-length Notch4, which is expectedly insensitive to GSI treatment.

**Fig. 1:**
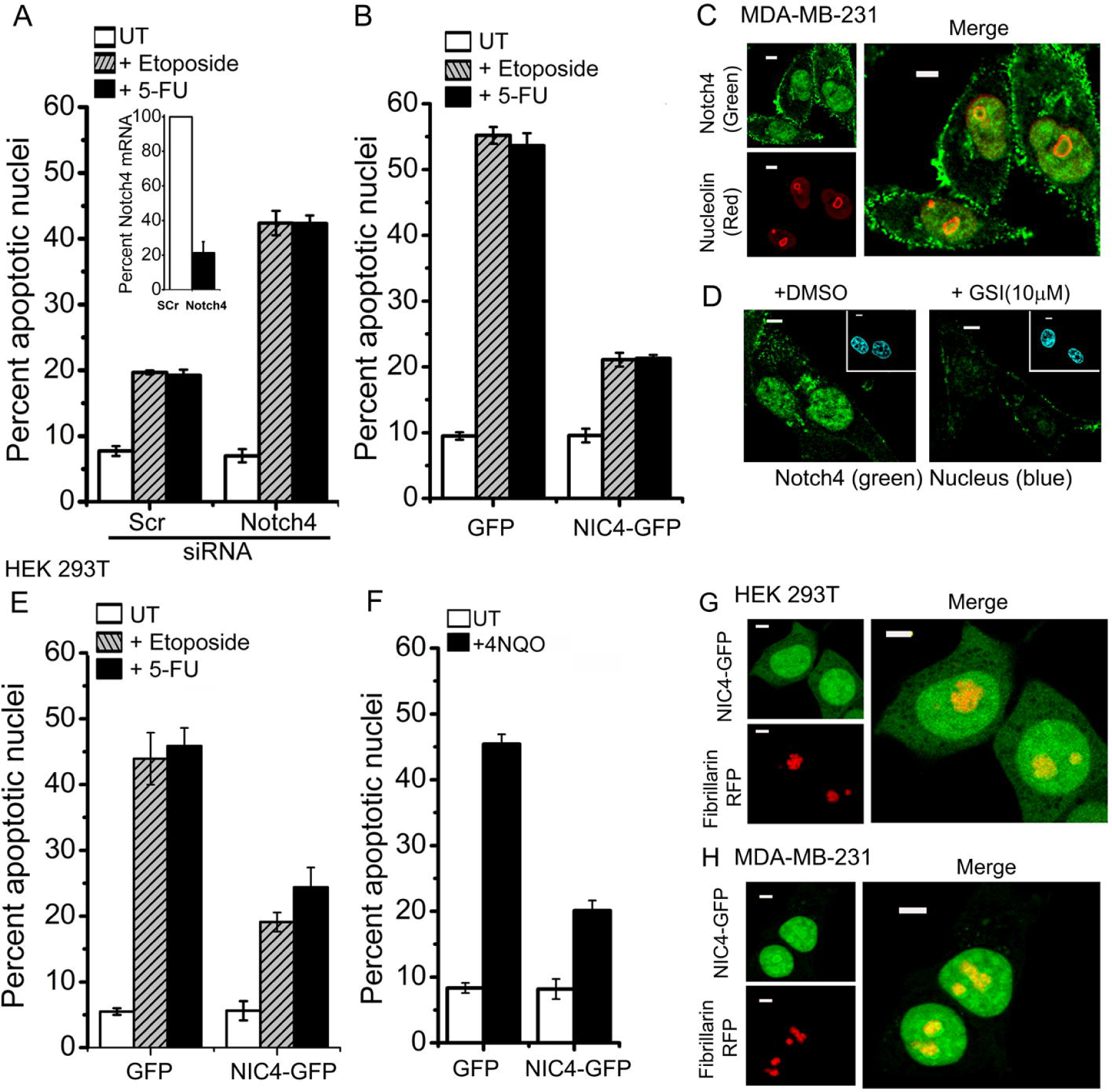
Notch4 signaling protects from apoptosis triggered by genotoxic agents. **A**, Induction of apoptotic nuclear damage in MDA-MB-231 cells pretreated with siRNA to Notch4 or a scrambled control for 24 h and then continued untreated (UT) or treated with etoposide (10μM) or 5-FU (10μM) for another 24 h in serum free medium. The inset shows percent Notch4 mRNA levels in scrambled and Notch4 siRNA transfected cells. **B**, Induction of apoptotic nuclear damage in MDA-MB-231 cells expressing the indicated plasmids and treated with etoposide or 5-FU for 48 h as in A. **C**, Representative confocal images of MDA-MB-231 cells stained for endogenous Notch4 (green) and Nucleolin (red) as described in methods (Manders Correlation Coefficient: 0.73 ± 0.23). **D**, Representative confocal images of MDA-MB-231 cells stained with the antibody to Notch4 in cells pre-treated with vehicle control or GSI-X (10μM) for 24 h in serum free medium. **E-F**, Percent apoptotic nuclear damage in HEK cells expressing GFP or NIC4-GFP and treated with etoposide or 5-FU (E) or 4NQO (5μM) (F) and cultured for 48 h in serum free medium. **G**, Representative confocal images of HEK cells co-expressing NIC4-GFP and Fibrillarin-RFP imaged 24h after transfection (Manders Correlation Coefficient: 0.87 ± 0.15). **H**, Representative confocal images of MDA-MB-231 cells co-expressing NIC4-GFP and Fibrillarin-RFP imaged 24 h after transfection (Manders Correlation Coefficient: 0.96 ± 0.07). In A, B, E and F, cells were stained with Hoechst 33342 to visualize nuclei. Data represent the mean ± S.D. of three independent experiments. Scale bar: 5μm

NIC4-mediated protection from genotoxic agents was also observed in the HEK cell line treated with etoposide or 5-FU (Fig. 1E) or following treatment with the quinolone compound 4-nitroquinoline *N*-oxide (4NQO) (Fig. 1F), which mimics radiation induced DNA damage [25]. Further, in HEK cells, NIC4-GFP was observed in the nucleoplasm as well as the nucleolus where it localized with co-transfected Fibrillarin-RFP, which marks nucleoli (Fig. 1G and Supplementary Figure 1C). Further, this pattern was also reproduced in MDA-MB-231 cells overexpressing NIC4-GFP, with the recombinant protein showing a distribution similar to endogenous NIC4 (Fig. 1H and Supplementary Figure 1D). Together these experiments establish that endogenous and overexpressed NIC4 is present in the nucleolus and nucleoplasm. Further, the experiments suggest that NIC4 regulates protection from genomic damage. Molecular complexes that coordinate the cellular response to DNA damage are present in the nucleolus [26,27]. Before assessing the functional consequences of its sub-cellular localization, we tested NIC4 dependence on nucleolar proteins for its anti-apoptotic activity.

### Dependence on nucleolar proteins for Notch4-mediated anti-apoptotic activity

NIC4-medaited anti-apoptotic activity was assessed following siRNA mediated ablation of nucleolar proteins, Nucleolin (NCL) or Fibrillarin (FBL), in HEK cells. Reduction in Nucleolin levels (Fig. 2A inset), abrogated NIC4-mediated protection from apoptosis triggered by etoposide or 5-FU (Fig. 2A). Similarly, ablation of Fibrillarin attenuated NIC4-mediated inhibition of etoposide or 4NQO induced apoptosis (Fig. 2B and 2C). Notably, when tested in same assays, Bcl-xL (the BCl-2 family anti-apoptotic protein) mediated protection from genomic damage was independent of FBL (Fig. 2B), indicating that ablation of NCL or FBL did not result in global changes and a specificity in interactions with NIC4. Further, NIC4-mediated anti-apoptotic activity was independent of its canonical nuclear partner RBPj-κ (Supplementary Figure 2A), which regulates Notch transcription. Together, these experiments demonstrate the activation of a distinct signaling cascade by NIC4. Next, we asked if canonical intermediates of the cellular DNA damage response machinery are required for NIC4 mediated anti-apoptotic activity.

**Fig. 2:**
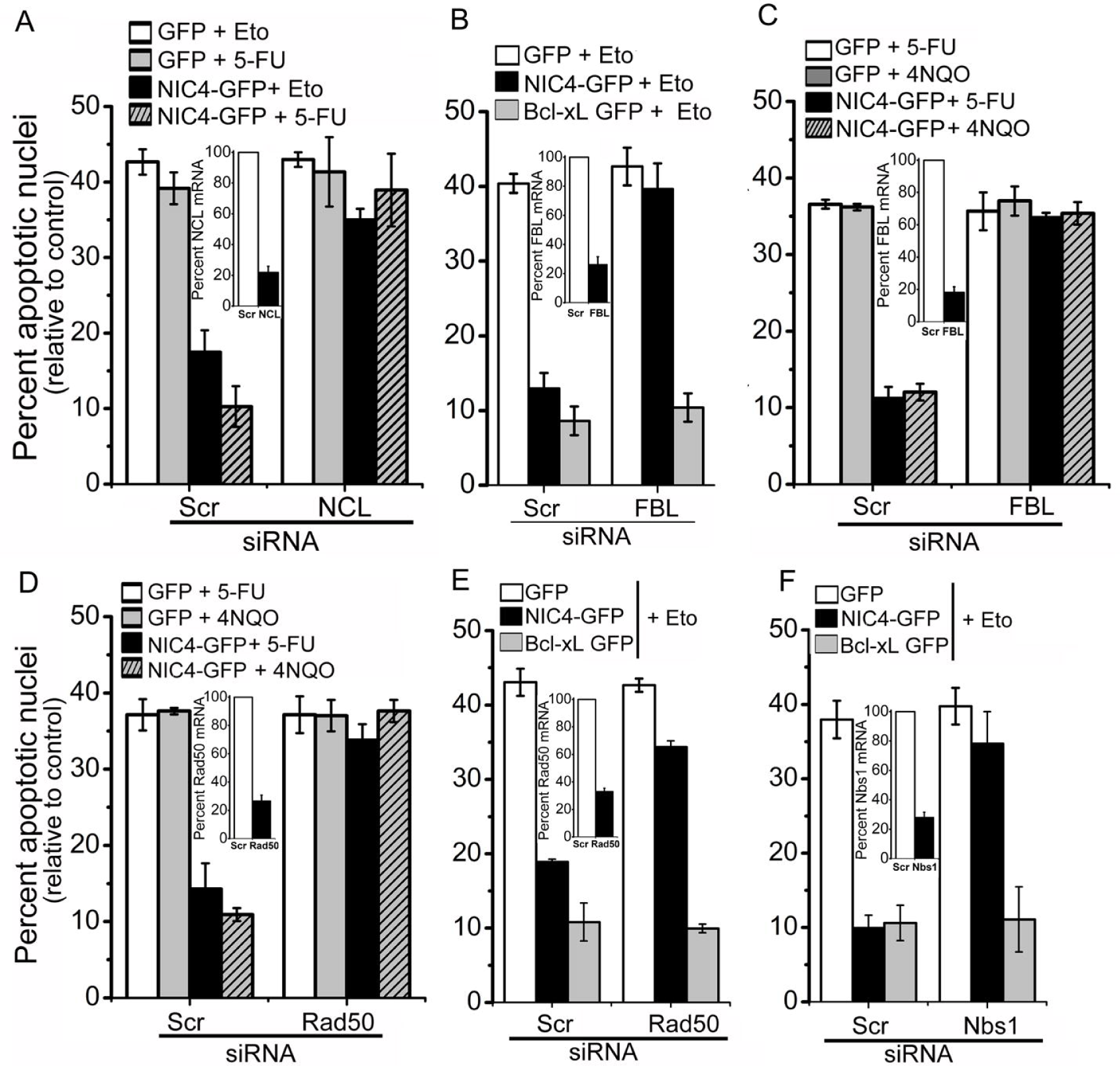
NIC4-mediated anti-apoptotic activity requires nucleolar proteins. **A-C**, Percent apoptotic nuclear damage in HEK cells expressing the indicated plasmids treated with etoposide (10μM), 5-FU (10μM) or 4NQO (5μM) for 48 h in serum-free medium, following treatment with siRNA to NCL (A) or FBL (B and C) or scrambled control. **D-E**, Percent apoptotic nuclear damage in HEK cells expressing the indicated plasmids treated with 5-FU (10μM) or 4NQO (5μM) (d) or etoposide (10μM) (E) for 48 h, following treatment with siRNA to Rad50 or scrambled control. **F**, Percent apoptotic nuclear damage in HEK cells expressing the indicated plasmids treated with etoposide in serum-free medium, following treatment with siRNA to Nbs1 or scrambled control. Insets in all panels show percent mRNA levels in scrambled and siRNA transfected cells. Data plotted are mean ± S.D. of three independent experiments.

### NIC4 mediated protection converges on cellular DNA damage-sensing proteins

The MRN (Mre11/Rad50/Nbs1) sensor complex comprises three proteins Mre11-Rad50-Nbs1 that act in concert to sense DNA damage and initiate repair [29,30,31]. Employing siRNA mediated ablations, we tested the requirement for Nbs1 and Rad50 for NIC4-mediated protection from genomic damage. Depletion of Rad50 abrogated NIC4-mediated inhibition of 4NQO, 5-FU or etoposide induced apoptosis (Fig. 2D and E), whereas Bcl-xL continued to inhibit apoptosis, following ablation of Rad50 (Fig. 2E). Similarly, while Nbs1 is required for NIC4 mediated activity, Bcl-xL mediated protection is independent of this intermediate (Fig. 2F). Thus, NIC4 signaling integrates with molecular complexes implicated in the sensing and repair of genomic damage, as well as proteins resident in the nucleolus – FBL and NCL - for anti-apoptotic activity.

### Dynamics of NIC4 localization to the nucleolus

The experiments reveal a hitherto undescribed aspect of Notch4 signaling vis-à-vis interactions with nucleolar proteins. In order to characterize the dynamics of the nuclear pools of NIC4, we employed <u>F</u>luorescence recovery <u>a</u>fter <u>p</u>hoto-bleaching (FRAP) analysis in live cells co-expressing NIC4-GFP and Fibrillarin-RFP, which indicated nucleoli. In this analysis, following a bleach of one spot marking nucleolar-localized NIC4-GFP, the recovery of GFP fluorescence was rapid with approximately 60% of the original intensity restored within a few seconds’ post photo-bleaching (Fig. 3A, 3B and Supplementary Figure 3A). Hence, while a large proportion of NIC4-GFP moves freely between the nucleoplasm and the nucleolus, a fraction of nucleolar NIC4-GFP has restricted mobility or is immobile. Expectedly, the recovery of the nucleolar protein Fibrillarin-RFP fluorescence following photo bleaching is low (Fig. 3A, 3B and Supplementary Figure 3A). NIC4-GFP dynamics remained the same in cells treated with etoposide (Fig. 3C and Supplementary Figure 3B).

**Fig. 3:**
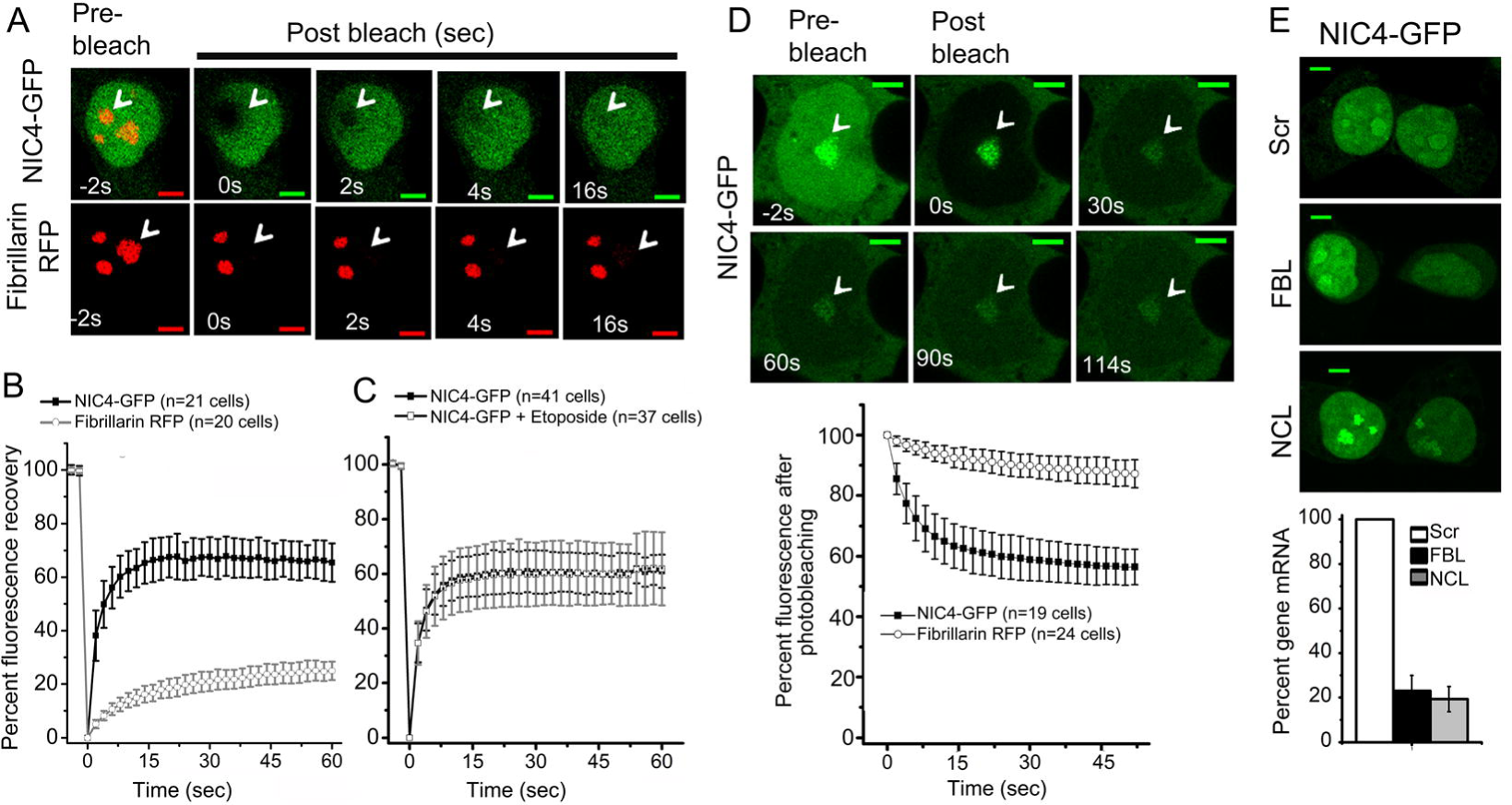
NIC4-GFP mobility assessed by FRAP and FLIP. **A**, Representative confocal images of a single cell co-expressing Fibrillarin-RFP and NIC4-GFP traced over time in FRAP analysis, bleached nucleolus is marked by the white arrowhead. **B-C**, Fluorescence intensity recovered over time plotted in the graph after photo bleaching in cells co-expressing Fibrillarin-RFP and NIC4-GFP (B) and in cells co-expressing Fibrillarin-RFP and NIC4-GFP cultured for 6 h with etoposide (10μM) in serum free medium (C). Data plotted are mean ± S.D. of a minimum of 20 cells in b and 37 cells in C. **D**, Loss of fluorescence intensity over time (FLIP) in cells co-expressing Fibrillarin-RFP and NIC4-GFP. The panel shows images of a single cell tracked over time, with the visualization spot (not bleached) marked by an arrow-head, surrounded by the bleached (dark) region over the duration of analysis plotted in the graph below. Data plotted are mean ± S.D. of a minimum of 19 cells in each condition. **E**, Representative confocal images of HEK cells expressing NIC4-GFP following treatment with siRNA to scrambled control (upper panel); FBL (middle panel) and NCL (lower panel). mRNA levels of indicated transcripts in scrambled and siRNA transfected cells is plotted. Images are representative of 30 cells in each condition. Scale bar: 5μm

The mobility of the nucleolar pools of NIC4-GFP was next estimated by <u>F</u>luorescence loss in <u>p</u>hoto-bleaching (FLIP) analysis in live cells. In this assay, a region of the nucleoplasm (which excludes the nucleolus) was photo-bleached and changes in fluorescence, if any, in the nucleolus tracked over time. If NIC4 is freely diffusible, diminished fluorescence in the nucleolus will be observed following the bleach of the surrounding region. The loss of fluorescence of Fibrillarin-RFP was expected to be minimal, as this protein was not detected in regions outside the nucleolus (Fig. 3D and Supplementary Figure 3C). NIC4-GFP fluorescence in the nucleolus is reduced by ∼40%, relative to the intensity at the onset of the assay and did not diminish further with time (Fig. 3D and inset images and Supplementary Figure 3C). This is consistent with the immobile fraction of nucleolar NIC4-GFP revealed by the FRAP analysis and raises the possibility of modifications or associations with nucleolar resident proteins that regulate NIC4-GFP dynamics. Notably, nucleolar localization of NIC4 was unchanged in cells ablated for either NCL or FBL (Fig. 3E and Supplementary Figure 3D).

### Nucleolar localization of Notch4 in breast cancer cell lines

To confirm that the results were not restricted to one cell line, the localization of endogenous Notch4 was assessed in HCC1086, BT-549, Hs578T, SUM149 and MCF-7 breast cancer cell lines. Notch4 (NIC4) staining was detected in the nucleoplasm and co-localized with Nucleolin in all cell lines examined (Fig. 4A, 4B and Supplementary Figure 4A-E). This is in striking contrast and distinct from the distribution of the closely related protein, Notch1, wherein the processed receptor is also nuclear localized but was excluded from the nucleolus (Fig. 4C and Supplementary Figure 4F-I). As seen in the MDA-MB-231 cells, siRNA mediated silencing of Notch4 increased sensitivity of Hs578T cells to apoptotic damage by etoposide or 5-FU (Fig. 4D), suggesting that Notch4 signaling sets the threshold for sensitivity to apoptotic damage. Ablation of Notch1 was without effect in this context (Fig. 4D).

**Fig. 4:**
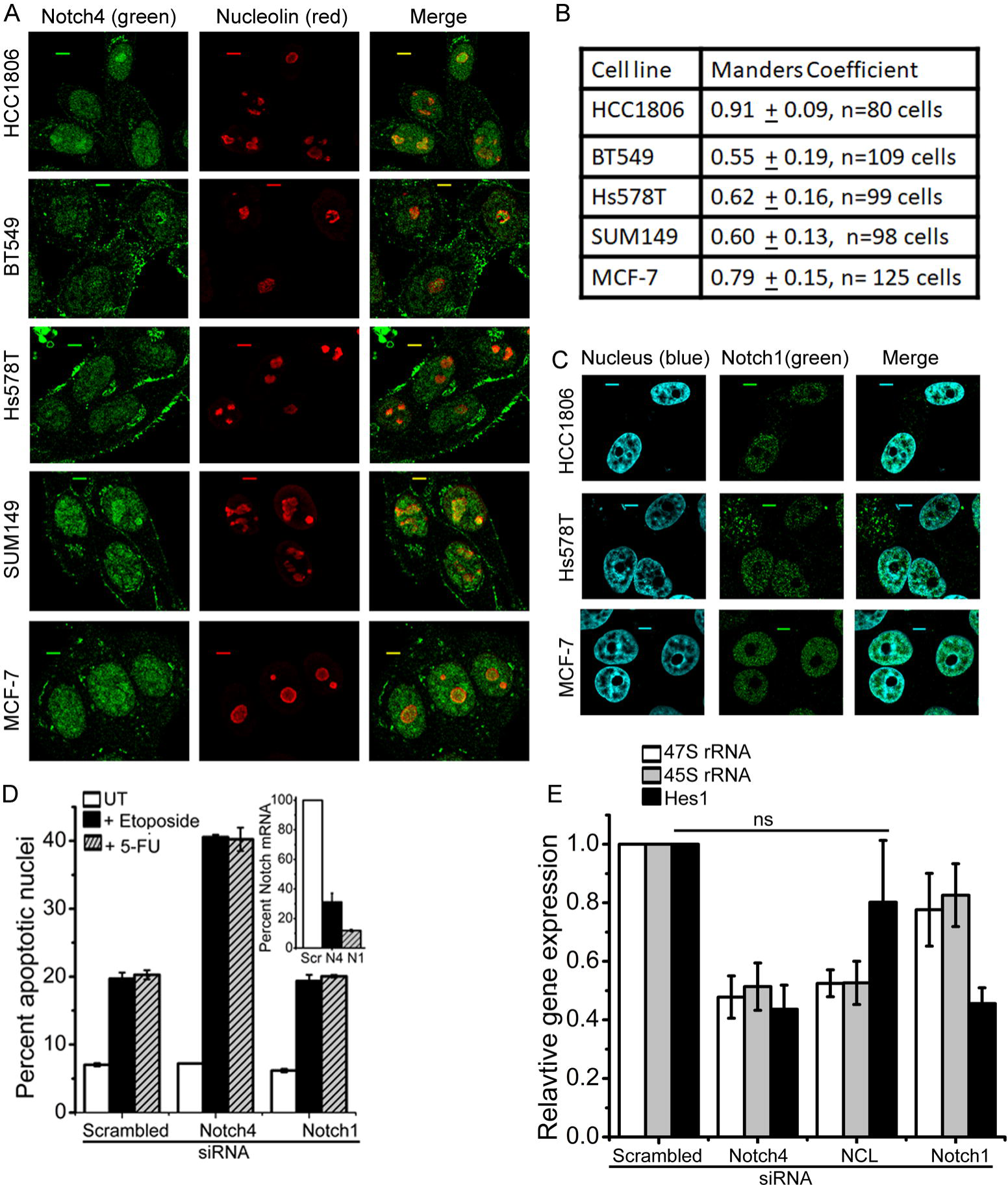
Notch4 localisation in breast cancer cell lines. **A**, Representative confocal images of indicated cell lines stained for endogenous Notch4 (green) or Nucleolin (red) as described in methods. Scale bar: 5μm **B**, Co-localisation of Nucleolin (red) with Notch4 (green) in indicated cell lines quantified by Manders Coefficient as described in methods and shown as mean ± S.D. **C**, Representative confocal images of some of the cell lines in **A**, stained for endogenous Notch1 (mN1A, green) and counterstained with Hoechst 33342 as described in methods. Scale bar: 5μm **D**, Induction of apoptotic nuclear damage in Hs578T cells pretreated with siRNA to Notch4 or Notch1 or scrambled control for 24 h and then cultured untreated (UT) or treated with etoposide (10μM) for another 24 h in serum free medium. Inset shows percent mRNA levels in scrambled control and siRNA transfected cells. **E**, mRNA levels of indicated genes in MDA-MB-231 cells treated with siRNA to Notch4 or Notch1 or NCL or scrambled control for 48 h. Data show the mean ± S.D. of two independent experiments in D and three independent experiments in E. ns: not significant, p > 0.05 (Student’s t-test).

The nucleolus is a hub for ribosome biogenesis and several nucleolar proteins are implicated in rRNA synthesis, maturation, and assembly [32,33,34]. Genome-wide screens have shown that nucleosome organization and ribosome biogenesis can be regulated by diverse cellular pathways [35,36], including signaling by a closely related protein Notch2 [37]. To test if nucleolar localization resulted in Notch4 regulation of ribosome synthesis, levels of 47s and 45s ribosomal RNA precursor transcripts were assessed, following the perturbation of Notch4. NCL ablation was included as a positive control in this experiment. We find that siRNA mediated ablation of Notch4 or Nucleolin, abrogated the induction of ribosomal 47S and 45S pre-rRNA transcripts to a similar extent (Fig. 4E), whereas the ablation of Notch1 was without effect (Fig. 4E). On the other hand, transcripts of the canonical Notch family target gene Hes1, were dramatically reduced following the ablation Notch4 or Notch1 (Fig. 4E). Despite a reduction in the levels of Hes1 transcripts in the NCL siRNA treated groups, relative to the scrambled control, the difference was not significant (Fig. 4E).

### Nucleolar localization is critical for NIC4-mediated anti-apoptotic activity

In the experiments that follow the analysis was extended to include spatially restricted NIC4 mutants to more directly ascertain functional consequences of nucleolar localization. We generated a recombinant form of NIC4 modified to increase residence time in the nucleolus by an additional nucleolar localization sequence (NoLS) derived from the NF-κB inducing kinase [38]. In order to disrupt NIC4 nucleolar localization [39], a second construct was generated with the positively charged Arginine residues 1490,1492 and 1501 – in NIC4 NoLS – replaced by the neutral amino acid Alanine (Fig. 5A). The sub-cellular localization of these constructs – NIC4-NoLS and NIC4-3RA respectively - was visualized by confocal microscopy following overexpression in HEK cells. The NIC4-3RA construct localized to the nucleus but, is excluded from the nucleolus (Fig. 5A and Supplementary Figure 5A). The NIC4-NoLS construct was expectedly enriched in the nucleolus but also detected at low levels in the nucleoplasm (Fig. 5A and Supplementary Figure 5B). Protection from genomic damage was abrogated in cells expressing the NIC4-3RA construct (Fig. 5B and 5C). However, protection from Thapsigargin (an ER stressor) induced apoptosis was comparable in NIC4 and NIC4-3RA expressing cells (Supplementary Fig. 5C), indicating that nucleolar localization was necessary for NIC4-mediated protection from genomic damage. On the other hand, inhibition of genomic damage by NIC4-NoLS was comparable to NIC4 (Fig. 5D).

**Fig. 5:**
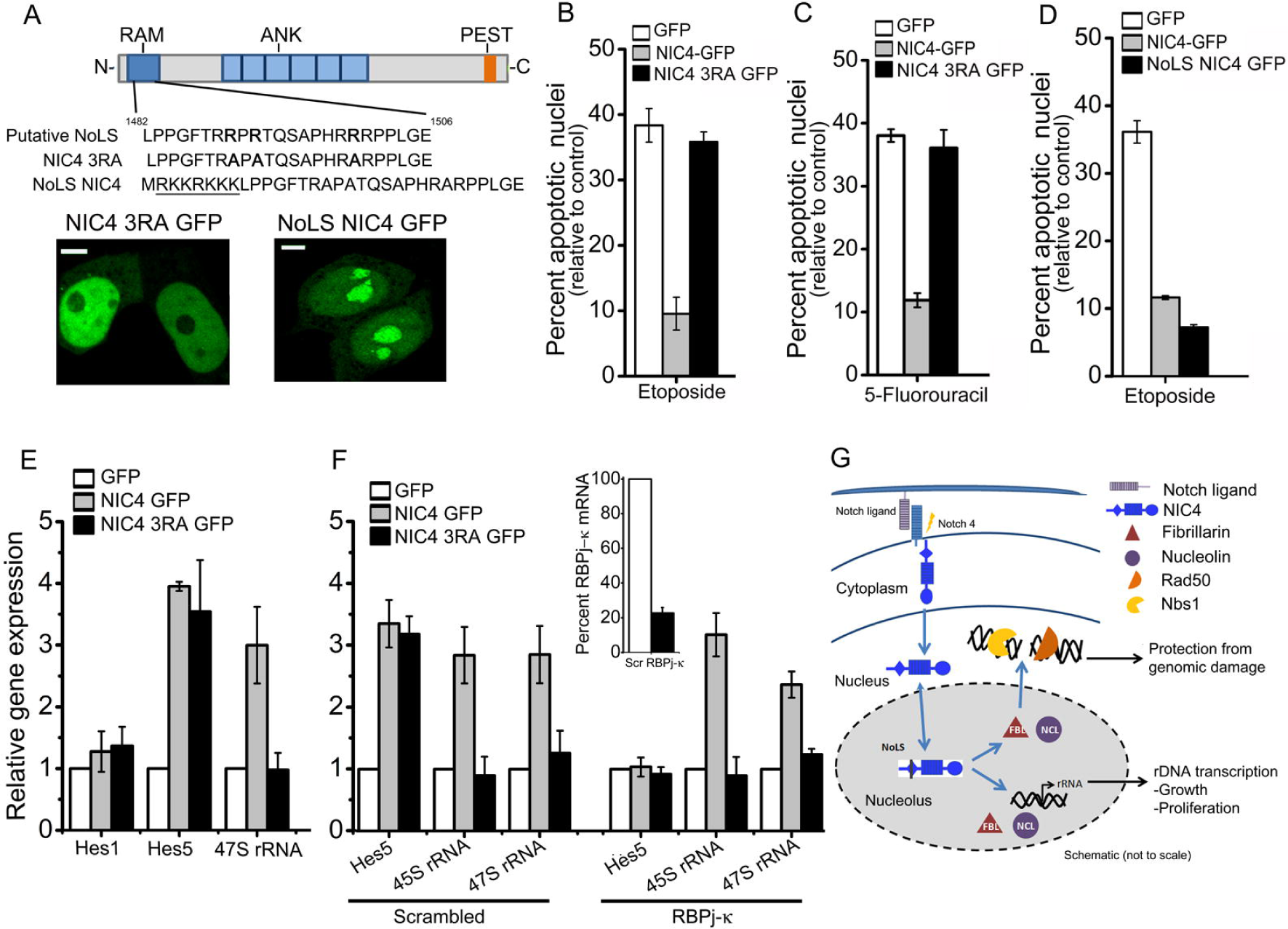
Nucleolar localization of NIC4 controls anti-apoptotic activity. **A**, Schematic of the putative NoLS sequence in NIC4; NIC4-3RA, with three arginine residues replaced by alanine to perturb the NoLS; and the additional NoLS at the NIC4 N-terminal; inset: Representative confocal images of HEK cells expressing NIC4-3RA GFP (left) and NoLS NIC4 GFP (right). Scale bar: 5μm. **B-C**, Induction of apoptotic nuclear damage in cells expressing GFP, NIC4-GFP or NIC43RA-GFP, treated with etoposide (10μM) (B) or 5-FU (10μM) (C) for 48h in serum free medium. **D**, Percent apoptotic nuclear damage in cells expressing GFP, NIC4-GFP or NoLS-NIC4 GFP, treated with etoposide for 48 h in serum free medium. **E**, Relative transcript levels of Hes1, Hes5 and 47S rRNA in cells transfected with indicated plasmids and cultured for 36 h in complete medium. **F**, Relative transcript levels of indicated genes in cells transfected with GFP, NIC4-GFP or NIC43RA-GFP and cultured for 36 h in complete medium following treatment with siRNA to RBPj-κ or NCL or scrambled control. Inset shows percent RBPj-κ mRNA levels in scrambled and siRNA transfected cells. Data plotted are mean ± S.D. of three independent experiments. **G**, The schematic (not drawn to scale) summarizes key observations of the current study. We show that NIC4 localizes to the nucleolus and demonstrate dependency on nucleolar proteins – Nucleolin and Fibrillarin – for protection from genotoxic agents. Further, NIC4 mediated anti-apoptotic activity is dependent on canonical regulators of the DNA damage response. The experiments reveal a functional role for a Nucleolar localization sequence (NoLS) in NIC4, which agrees with NIC4 regulation of pre-ribosomal RNA transcription reported in this study.

We next tested if NIC4-mediated rDNA transcription was detected in cells expressing NIC4-GFP or the spatially modified NIC43RA-GFP recombinant. Compared to the GFP transfected group, an increase in pre-rRNA 47s and 45s transcripts was observed in cells expressing NIC4, but not in cells expressing NIC4-3RA (Fig. 5E). The induction of Hes5 transcripts was comparable in both groups (Fig. 5E), confirming that activation of transcriptional outcomes by NIC4-3RA was not entirely abrogated. Hes5 induction was reduced in cells expressing NIC4-NoLS (Supplementary Figure 5D), indicating differential outputs associated with NIC4 localization. The transcription factor RBPj-κ is a key intermediate in the transcription of Notch (canonical) target genes. Confirming this dependence, Hes5 transcripts were not induced when NIC4 or NIC4-3RA were expressed in cells treated with siRNA to RBPj-κ (Fig. 5F). However, the induction of 47S and 45S rRNA transcripts was not compromised by the ablation of RBPj-κ and remained comparable to the control siRNA (scrambled) group (Fig. 5F). This is consistent with differential regulation of transcription occurring in the nucleolus and nucleoplasm. Together, these experiments illustrate that spatial segregation of Notch4 results in diverse signaling outcomes that are distinct and uncoupled from the canonical pathway activated by NIC4 signaling in the nucleoplasm (Fig. 5G).

## Discussion

Notch family proteins integrate with a range of signaling cascades that underlie pleiotropic outcomes of Notch signaling. In this study we report new aspects of Notch 4 signaling, which show that spatial regulation of Notch4 results in the diversification of signaling pathways initiated by Notch4 activity. To our knowledge, this is the first demonstration of the nucleolar localization of Notch4 and functional consequences of this sub-cellular distribution. Both protection from genomic damage as well as the regulation of ribosome synthesis was compromised following disruption of the NoLS in NIC4. Nucleolar localization of NIC4 was established for endogenous and over-expressed protein, using imaging analysis also coupled to biophysical measurements. NIC4 mediated protection from genomic damage was independent of canonical signaling through RBPj-κ, a feature shared with the closely related protein Notch1 [28]. NIC4 localization and regulation of ribosome biogenesis was validated in multiple cell lines as well as in cells overexpressing NIC4 and spatially restricted forms of the same.

We demonstrate dependence on nucleolar proteins Nucleolin and Fibrillarin for Notch4-mediated protection from genomic damage. This dependency is specific to Notch4 as it is not observed with Bcl-xL, a member of the Bcl-2 family, which also protects from apoptosis triggered by genotoxic agents. We show that ablation of Notch4 but not Notch1 in the breast cancer cell lines MDA-MB-231 and Hs578T modulated sensitivity to genotoxic stress. These experiments and observations with modified forms of NIC4, provide compelling evidence that NIC4 signaling integrates with proteins in the nucleolus to mediate a distinct signaling pathway for the protection from DNA damage. While the sub-cellular distribution of NIC4 in the nucleolus is not regulated by Fibrillarin or Nucleolin, these proteins are key, non-redundant intermediates of the NIC4-activated signaling cascade. Somewhat unexpectedly we observed that NIC4 controls transcription of ribosomal pre-rRNA, implicating Notch4 in ribosome biogenesis, although the physiological consequences of this central cellular process, remains to be investigated. Together, these observations suggest an additional layer of regulation of Notch4 activity with implications for growth regulation in development and disease [40, 41, 42, 43].

This study raises broader questions of the regulation of Notch family proteins. Notch4 is upregulated in breast cancers and its inhibition reported to reduce invasiveness and tumorigenic properties in models of breast cancer and in cell lines [44,45,46,47]. Evidence that specific increases in Notch4, is coincident with reductions in other receptors indicating a switch to the activation of Notch4 signaling in cancer stem cells, may correlate with the acquisition of drug-resistance in breast cancers [46,48,49,50]. The inhibition of etoposide or 5-FU induced apoptosis in cells expressing NIC4 is consistent with reports that Notch inhibition sensitizes various tumors to chemotherapy [51]. To this end, our experiments have identified previously unappreciated interactions arising from its nucleolar location that are critical to Notch4 signaling. Collectively, this study provides yet another example of how spatial regulation of the Notch family [14,16,17,52,53] expands the repertoire of signaling pathways activated by these receptors.

## Supporting information

Saini_Sarin Suppl file

## Non-Standard Abbreviations

4NQO: 4-Nitroquinoline*N*-oxide,
5-FU: 5 Fluorouracil;
Bcl-xL: B-cell lymphoma extra-large;
FBL: Fibrillarin;
Hes1: hairy and enhancer of split-1;
NIC: Notch intracellular domain;;
NCL: Nucleolin;
Nbs1: Nijmegen Breakage Syndrome 1;
RBPj-κ: recombination signal – binding protein-Jκ.

## Acknowledgements

We acknowledge the Central Imaging and Flow Cytometry Facility (CIFF) at NCBS and InStem, Bangalore. This work was supported by a grant from the Science and Engineering Research Board (EMR/2015/001023), Department of Science and Technology (DST), Govt. of India and core funds from the Institute for Stem Cell Science and Regenerative Medicine (inStem), Bellary Road, Bangalore, India to AS.

## Author Contributions

NS performed and analyzed experiments and contributed to the writing of the manuscript; AS contributed to experiment design, analysis and writing of the manuscript.

## References

1. Nwabo Kamdje AH, Mosna F, Bifari F, Lisi V, Bassi G, Malpeli G et al. Notch-3 and Notch-4 signaling rescue from apoptosis human B-ALL cells in contact with human bone marrow-derived mesenchymal stromal cells. Blood 2011; 118: 380–390.

2. Perumalsamy LR, Nagala M, Sarin A. Notch-activated signaling cascade interacts with mitochondrial remodeling proteins to regulate cell survival. Proc Nat Acad Sci 2010; 107: 6882–6887.

3. MacKenzie F, Duriez P, Wong F, Noseda M, Karsan A. Notch4 inhibits endothelial apoptosis via RBPjk-dependent and -independent pathways. J Biol Chem 2004; 279: 11657–11663.

4. Marcel N, Sarin A. Notch1 regulated autophagy controls survival and suppressor activity of activated murine T-regulatory cells. eLIFE 2016; 5: e14023.

5. Dumortier A, Wilson A, MacDonald HR, Radtke F. Paradigms of Notch signaling in mammals. Int J Hematol 2005; 82: 277–284.

6. Kopan R, Ilagan MXG. The Canonical Notch Signaling Pathway: Unfolding the Activation Mechanism. Cell 2009; 137: 216–233.

7. Gordon WR, Arnett KL, Blacklow S. The molecular logic of Notch signaling– a structural and biochemical perspective. J Cell Sci 2008; 121: 3109–3119.

8. Schroeter EH, Kisslinger JA, Kopan R. Notch-1 signalling requires ligand-induced proteolytic release of intracellular domain. Nature 1998; 393: 382–386.

9. Brou C, Logeat F, Gupta N, Bessia C, LeBail O, Doedens JR et al. A novel proteolytic cleavage involved in Notch signaling. Mol Cell 2000; 5: 207–216.

10. Mumm JS, Schroeter EH, Saxena MT, Griesemer A, Tian X, Pan DJ et al. A ligand-induced extracellular cleavage regulates γ-Secretase-like proteolytic activation of Notch1. Mol Cell 2000; 5: 197–206.

11. Jarriault S, Brou C, Logeat F, Schroeter EH, Kopan R, Israel A. Signalling downstream of activated mammalian Notch. Nature 1995; 377: 355–358.

12. Wu L, Sun T, Kobayashi K, Gao P, Griffin JD. Identification of a family of mastermind-like transcriptional coactivators for mammalian notch receptors. Mol Cell Biol 2002; 22: 7688–7700.

13. Hori K, Fostier M, Ito M, Fuwa TJ, Go MJ, Okano H et al. Drosophila Deltex mediates Suppressor of Hairless-independent and late-endosomal activation of Notch signaling. Development 2004; 131: 5527–5537.

14. Mukherjee T, Kim WS, Mandal L, Banerjee U. Interaction between Notch and Hif-a in development and survival of Drosophila blood cells. Science 2011; 1210: 1–5.

15. Lee KS, Wu Z, Song Y, Mitra SS, Feroze AH, Cheshier SH et al. Roles of PINK1, mTORC2, and mitochondria in preserving brain tumor-forming stem cells in a noncanonical Notch signaling pathway. Genes Dev 2013; 27: 2642–2647.

16. Hyun MS, Tilahun ME, Cho OH, Chandiran K, Kuksin CA, Keerthivasan S et al. Notch1 can initiate NF-κB activation via cytosolic interactions with components of the T cell signalosome. Front Immunol 2014; 5: 2–15.

17. Perumalsamy LR, Nagala M, Banerjee P, Sarin A. A hierarchical cascade activated by non-canonical Notch signaling and the mTOR–Rictor complex regulates neglect-induced death in mammalian cells. Cell Death Differ 2009; 16: 879–889.

18. Callahan R, Raafat A. Notch signaling in mammary gland tumorigenesis. J. Mammary Gland Biol. Neoplasia 2001; 6: 23–36.

19. Naik S, Macfarlane M, Sarin A. Notch4 signaling confers susceptibility to trail-induced apoptosis in breast cancer cells. J Cell Biochem 2015; 116: 1371–1380.

20. Fukusumi T, Guo TW, Sakai A, Ando M, Ren S, Haft S et al. The Notch4– Hey1 pathway induces Epithelial–Mesenchymal transition in Head and Neck Squamous Cell Carcinoma. Clin Cancer Res 2018; 24: 619–633.

21. Livak KJ, Schmittgen TD. Analysis of relative gene expression data using Real-Time Quantitative PCR and the 2-ΔΔCT method. Methods 2001; 25: 402–408.

22. Soubeyrand S, Pope L, Haché RJG. Topoisomerase IIα-dependent induction of a persistent DNA damage response in response to transient etoposide exposure. Mol Oncol 2010; 4: 38–51.

23. Houghton JA, Tillman DM, Harwood FG. Ratio of 2’-deoxyadenosine-5’triphosphate/Thymidine-5’-triphosphate influences the commitment of human colon carcinoma cells to Thymineless death. Clin Cancer Res 1995; 1: 723–730.

24. Tajrishi MM, Tuteja R, Tuteja N. The most abundant multifunctional phosphoprotein of nucleolus Nucleolin. Commun Integr Biol 2011; 4: 267–275.

25. Miao ZH, Rao VA, Agama K, Antony S, Kohn KW, Pommier Y. 4-Nitroquinoline-1-Oxide induces the formation of cellular Topoisomerase I-DNA cleavage complexes. Cancer Res 2006; 66: 6540–6546.

26. Kobayashi J, Fujimoto H, Sato J, Hayashi I, Burma S, Matsuura S et al. Nucleolin participates in DNA double-strand break induced damage response through MDC11. PLoS One 2012; 7: e49245.

27. Ogawa LM, Baserga SJ. Crosstalk between the nucleolus and the DNA damage response. Mol Biosyst 2018; 13: 443–455.

28. Vermezovic J, Adamowicz M, Santarpia L, Rustighi A, Forcato M, Lucano C et al. Notch is a direct negative regulator of the DNA-damage response. Nat Struct Mol Biol 2015; 22: 1–11.

29. Dinkelmann M, Spehalski E, Stoneham T, Buis J, Wu Y, Sekiguchi JM et al. Multiple functions of MRN in end-joining pathways during isotype class switching. Nat Struct Mol Biol 2009; 16: 808–813.

30. Symington LS, Gautier J. Double-Strand Break End Resection and Repair Pathway Choice. Annu Rev Genet 2011; 45: 247–271.

31. Taylor EM, Cecillon SM, Bonis A, Chapman JR, Povirk LF, Lindsay HD. The Mre11 / Rad50 / Nbs1 complex functions in resection-based DNA end joining in Xenopus laevis. Nucleic Acids Res 2010; 38: 441–454.

32. Rickards B, Flint SJ, Cole MD, Leroy G. Nucleolin is required for RNA polymerase I Transcription *in vivo*. Mol Cell Biol 2007; 27:937–948.

33. Murano K, Okuwaki M, Hisaoka M, Nagata K. Transcription regulation of the rRNA gene by a multifunctional nucleolar protein, B23/Nucleophosmin, through its histone chaperone activity. Mol Cell Biol 2008; 28: 3114–3126.

34. Bierhoff IA, Krogh N, Tessarz P, Ruppert T, Nielsen H, Grummt I. SIRT7-dependent deacetylation of Fibrillarin controls histone H2A methylation and rRNA synthesis during the cell cycle. Cell Rep 2018; 25: 2946–2954.

35. Farley-Barnes KI, McCann KL, Ogawa LM, Merkel J, Surovtseva YV, Baserga SJ. Diverse Regulators of Human Ribosome Biogenesis Discovered by Changes in Nucleolar Number. Cell Rep 2018; 13: 1923–1934.

36. Neumüller RA, Gross T, Samsonova AA, Vinayagam A, Buckner M, Founk K, et al. Conserved regulators of nucleolar size revealed by global phenotypic analyses. Sci Signal 2013;6: ra70.

37. Badertscher L, Wild T, Montellese C, Alexander LT, Bammert L, Sarazova M et al. Genome-wide RNAi screening identifies protein modules required for 40S subunit synthesis in human Cells. Cell Rep 2015; 13: 2879–2891.

38. Birbach A, Bailey ST, Ghosh S, Schmid JA. Cytosolic, nuclear and nucleolar localization signals determine subcellular distribution and activity of the NF-κB inducing kinase NIK. J Cell Sci 2004; 40: 3615–3624.

39. Scott MS, Boisvert FM, McDowall MD, Lamond AI, Barton GJ. Characterization and prediction of protein nucleolar localization sequences. Nucleic Acids Res 2010; 38: 7388–7399.

40. Susaki WK, Krogh N, Tessarz P, Ruppert T, Nielsen H, Grummt I. Biosynthesis of ribosomal RNA in nucleoli regulates pluripotency and differentiation ability of pluripotent stem cells. Stem Cells 2014; 12: 3099–3111.

41. Trainor PA, Merrill AE. Ribosome biogenesis in skeletal development and the pathogenesis of skeletal disorders. BBA-Mol Basis Dis 2014; 1842: 769–778.

42. Zhang C, Yin C, Wang L, Zhang S, Qian Y, Ma J et al. HSPC111 governs breast cancer growth by regulating ribosomal biogenesis. Mol Cancer Res 2014; 12: 583–595.

43. Belin S, Beghin A, Solano-Gonzàlez E, Bezin L, Brunet-Manquat S, Textoris J et al. Dysregulation of ribosome biogenesis and translational capacity is associated with tumor progression of human breast cancer cells. PLoS One 2009; 4: 1–13.

44. Gallahan D, Callahan R. The mouse mammary tumor associated gene INT3 is a unique member of the NOTCH gene family (NOTCH4). Oncogene 1997; 14, 1883–1890.

45. Nagamatsu I, Onishi H, Matsushita S, Kubo M, Kai M, Imaizumi A et al. NOTCH4 is a potential therapeutic target for triple-negative breast cancer. Anticancer Res 2014; 34: 69–80.

46. Harrison H, Farnie G, Howell SJ, Rock RE, Stylianou S, Brennan KR et al. Regulation of breast cancer stem cell activity by signaling through the Notch4 receptor. Cancer Res 2010; 70: 709–719.

47. D’Angelo RC, Ouzounova M, Davis A, Choi D1, Tchuenkam SM, Kim G et al. Notch reporter activity in breast cancer cell lines identifies a subset of cells with stem cell activity. Mol Cancer Ther 2015; 14: 779–787.

48. Lombardo Y, Faronato M, Filipovic A, Vircillo V, Magnani L, Coombes RC. Nicastrin and Notch4 drive endocrine therapy resistance and epithelial to mesenchymal transition in MCF7 breast cancer cells. Breast Cancer Res 2014; 16: R62.

49. Simões BM, O’Brien CS, Eyre R, Silva A, Yu L, Sarmiento-Castro A et al. Anti-estrogen resistance in human breast tumors is driven by Jag1-Notch 4-dependent cancer stem cell activity. Cell Rep 2015; 12: 1968–1977.

50. Qi XS, Pajonk F, Mccloskey S, Low DA, Kupelian P, Steinberg M. Radioresistance of the breast tumor is highly correlated to its level of cancer stem cell and its clinical implication for breast irradiation. Radiother Oncol 2017; 124: 455–461.

51. Schott AF, Landis MD, Dontu G, Griffith KA, Layman RM, Krop I et al. Preclinical and clinical studies of gamma secretase inhibitors with docetaxel on human breast tumors. Clin Cancer Res 2013; 19: 1512–1524.

52. Acosta H, López SL, Revinski DR, Carrasco AE. Notch destabilises maternal beta-catenin and restricts dorsal-anterior development in Xenopus. Development 2011; 138: 2567–2579.

53. Jin S, Mutvei AP, Chivukula IV, Andersson ER, Ramsköld D, Sandberg R et al. Non-canonical Notch signaling activates IL-6/JAK/STAT signaling in breast tumor cells and is controlled by p53 and IKKα/IKKβ. Oncogene 2013; 32: 4892–4902.

