## Supplementary material for "Signaling activated by nucleolar localised Notch4 Intracellular Domain underlies protection from genomic damage": Saini_Sarin Suppl file

**This file includes:**

Supplementary figures S1-S5

Supplementary figure legends S1-S5

**Supplementary figure 1.**

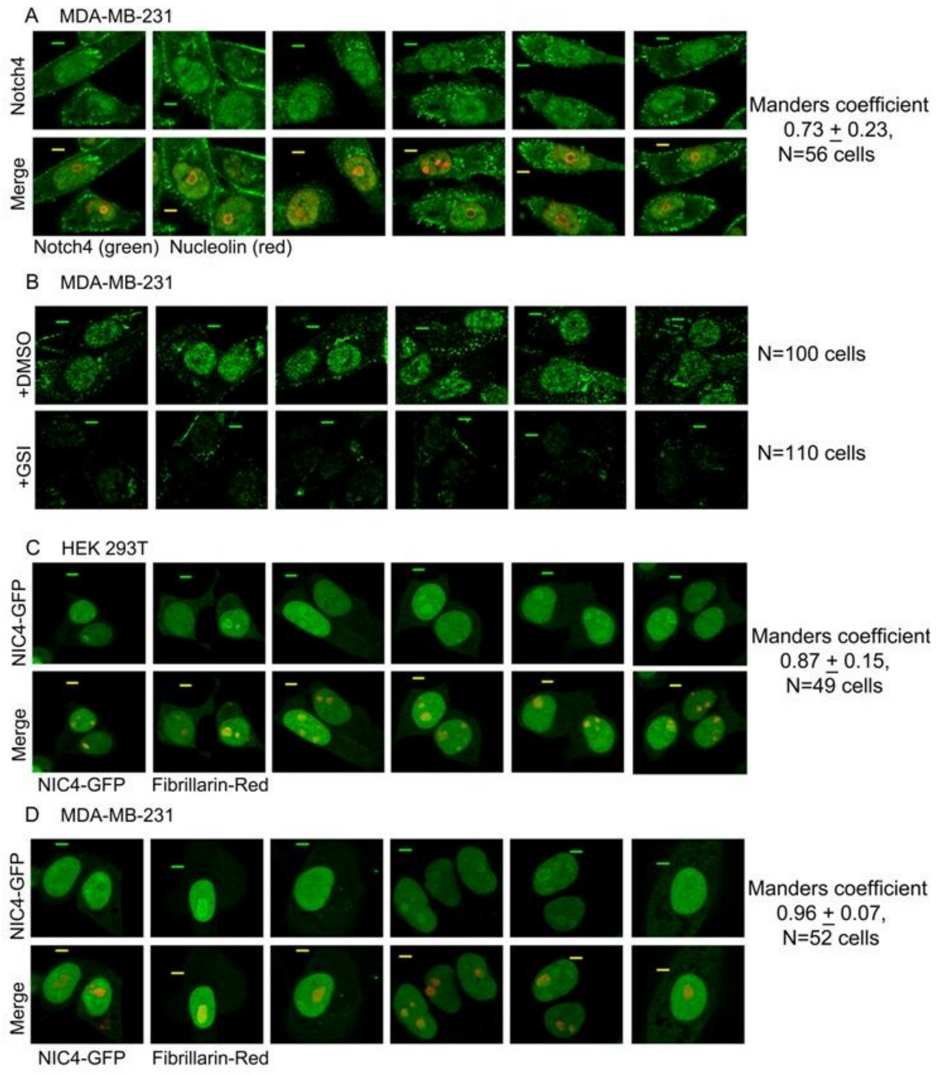


**Figure 1. Nucleolus localisation of NIC4**

**A,** Representative confocal images of MDA-MB-231 cells stained for endogenous Notch4 (green) and Nucleolin (red) as described in methods.

**B,** Representative confocal images of MDA-MB-231 cells treated with vehicle control or GSI-X (10μM) for 24 h in serum free medium were stained for Notch4 (green). Scale bar: 5μm.

**C,D** Representative confocal images of HEK (C) and MDA-MB-231 (D) cells co-expressing NIC4-GFP and Fibrillarin-RFP imaged 24 h after transfection.

In A,C,D co-localisation of Nucleolin (red) with Notch4 (green) was quantified by Manders coefficient as described in methods and shown as mean ± S.D.

**Supplementary figure 2.**

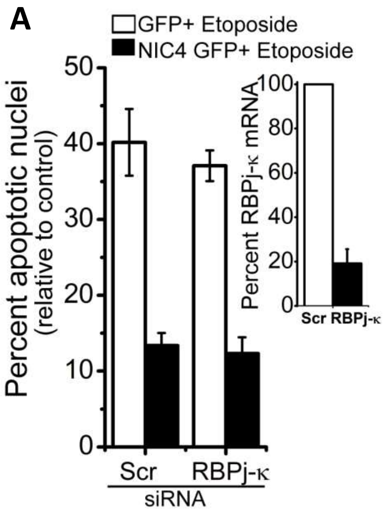


**Figure 2. NIC4 mediated anti-apoptotic activity does not require RBPj-κ**

**A** Percent apoptotic nuclear damage in HEK cells expressing GFP or NIC4-GFP treated with etoposide (10μM) for 48 h in serum-free medium, following treatment with siRNA to RBPj-κ or scrambled control; inset shows mRNA levels of RBPj-κ in cells treated with scrambled control or RBPj-κ siRNA. Data plotted are mean ± S.D. of three independent experiments.

**Supplementary figure 3.**

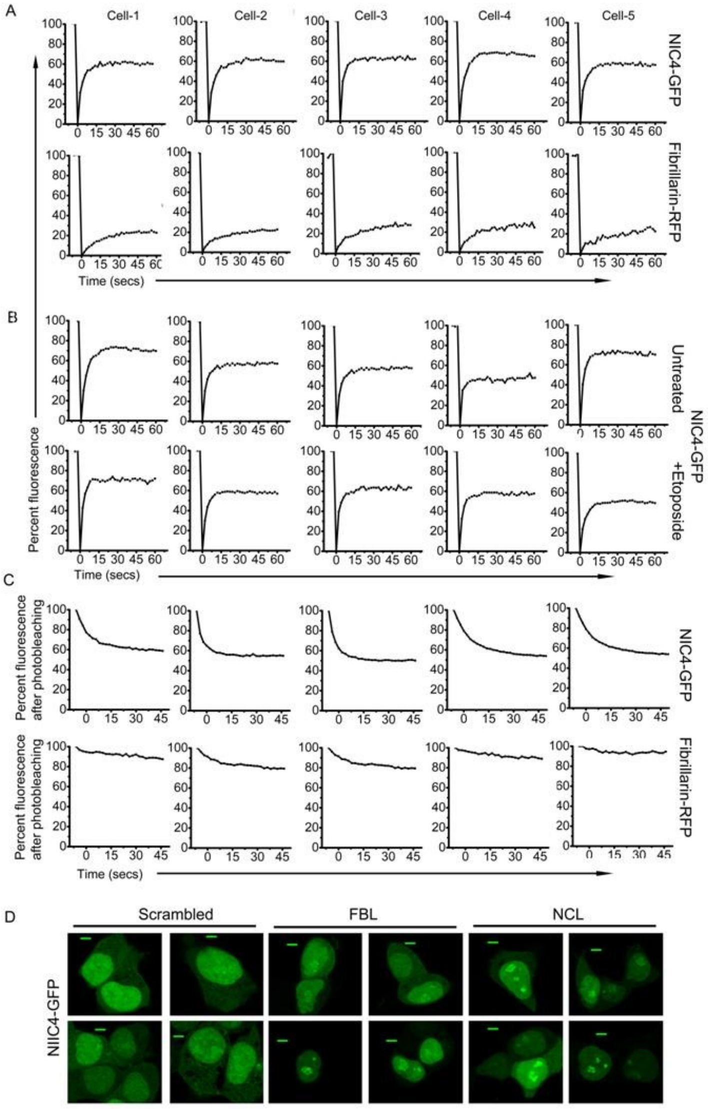


**Figure 3 NIC4-GFP has dynamic mobility between the nucleolus and nucleoplasm**

**A** Percent fluorescence intensity of NIC4-GFP (top) and Fibrillarin-RFP (bottom) recovered over time plotted in graph after photo-bleaching the nucleolus in cells co-expressing NIC4-GFP and Fibrillarin-RFP.

**B** Percent fluorescence intensity of NIC4-GFP recovered over time plotted in graph after photo-bleaching the nucleolus in cells cultured for 24 hours after transfection with NIC4-GFP and Fibrillarin-RFP, and left untreated (top) or treated (lower) with etoposide (10μM) for additional 6 h in serum free medium.

**C** Percent fluorescence intensity of NIC4-GFP (top) and Fibrillarin-RFP (lower) in the nucleolus after photo-bleaching the nucleoplasm in cells 24 hours after transfection with NIC4-GFP and Fibrillarin-RFP.

**D** Representative confocal images of HEK 293T cells expressing NIC4-GFP following transfection with scrambled control (left panel) or FBL (middle panel ) or NCL (right panel) siRNA (N>30 cells); Scale bar: 5μm.

**Supplementary figure 4.**

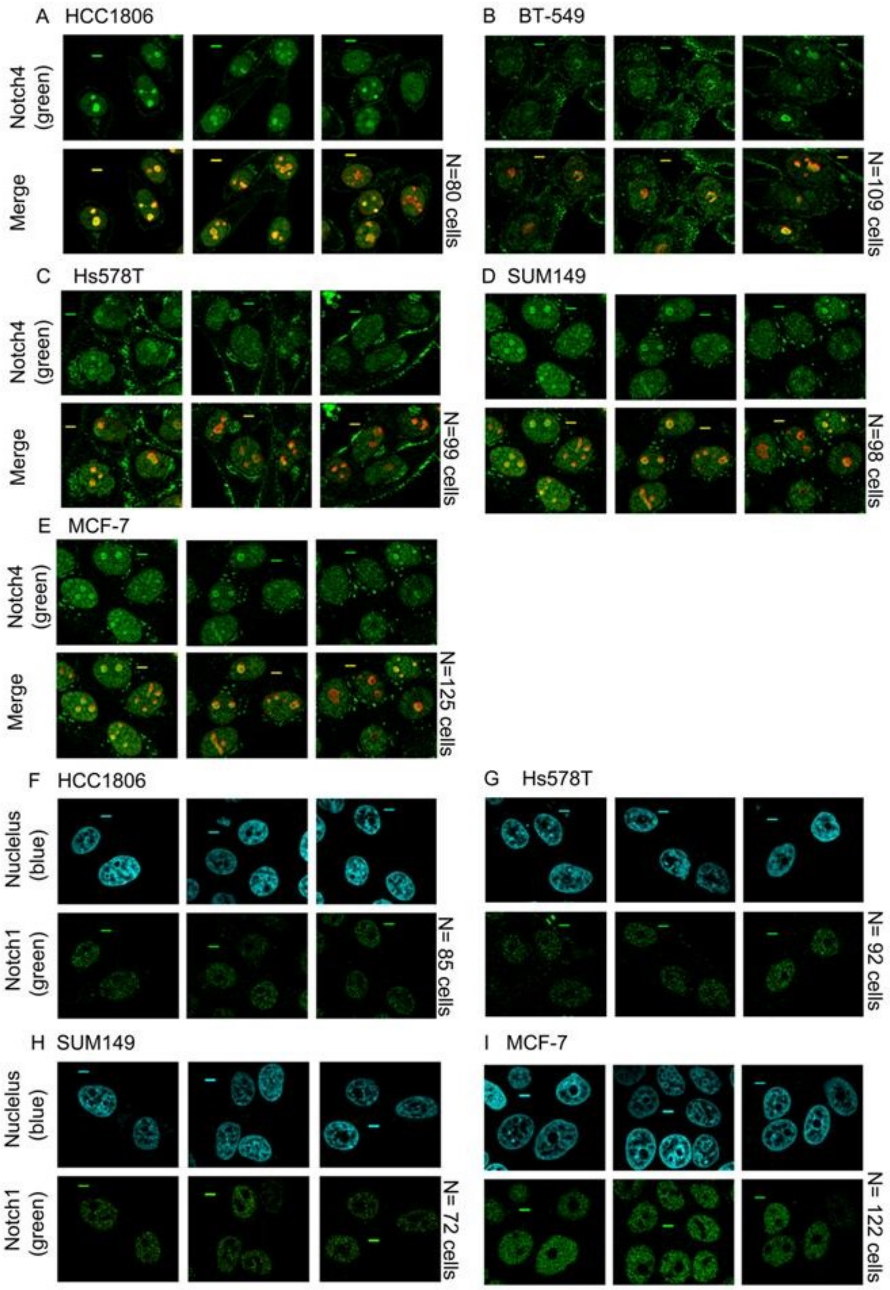


**Figure 4. Notch4 but not Notch1 localises to the nucleolus in breast cancer cell lines**

**A-E** Representative confocal images of HCC1806 (A), BT-549 (B), Hs578T (C), SUM149 (D) and MCF-7 (E) cell lines stained for endogenous Notch4 (green) and Nucleolin (red) as described in methods.

**F-I** Representative confocal images of HCC1806 (F), Hs578T (G), SUM149 (H) and MCF-7 (I) stained for Notch1 (mN1A, green) and counterstained with Hoechst 33342 (blue) as described in methods. Scale bar: 5µm.

**Supplementary figure 5.**

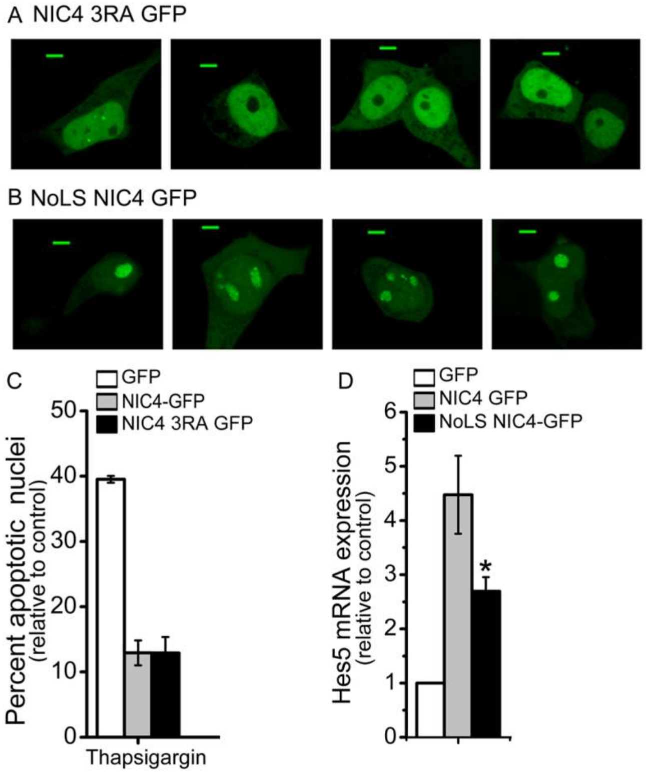


**Figure 5. Putative NoLS regulates nucleolar localisation of NIC4**

**A,B** Representative confocal images of HEK cells expressing NIC4 3RA GFP (n=25 cells) (A) or NoLS NIC4 GFP (n=25 cells) (B) imaged 24 h after transfection; Scale bar: 5μm.

**C** Induction of apoptotic nuclear damage in cells expressing GFP, NIC4-GFP or NIC43RA-GFP, treated with thapsigargin (10μM) for 24 h in serum free medium.

**D** Relative Hes5 mRNA levels in cells transfected with GFP, NIC4-GFP or NoLS NIC4 GFP, and cultured for 36 h in complete medium. * significant $p < 0.05$ (student's t-test)

Data plotted are mean ± S.D. of three independent experiments.
